# Circadian Regulation of Taste Sensitivity in Human and Rat Active Periods

**DOI:** 10.64898/2026.04.15.718832

**Authors:** Hiroko Mochizuki-Kawai, Tingbi Xiong, Shota Shimoda, Michimasa Toyoshima, Reina Tachihara, Hideaki Oike, Takayuki Kawai, Fumiyo Hayakawa, Yuko Nakano, Rei Osuga, Mio Kamei, Yuko Kusakabe, Kazuo Yamada

## Abstract

Most organisms exhibit endogenous 24-h rhythms that align physiology and behavior with the time of day; however, whether taste sensitivity oscillates in a circadian manner remains unclear, limiting our understanding of its role in feeding behavior. Here, we show that sensitivity to sweet and bitter tastes exhibits circadian modulation, peaking during the active period: daytime in humans and the dark phase in nocturnal rats. In humans, the rhythmicity of sweet taste was evident in detection but not recognition thresholds, indicating circadian modulation at the level of primary sensory processing rather than higher-order cognitive processes. In rats, increased sensitivity during the dark phase was accompanied by upregulation of taste-related genes in type II taste cells in tongue. These findings provide evidences that circadian regulation of taste sensitivity is conserved across species and appears to operate predominantly at the peripheral level. This mechanism may have evolved to optimize energy intake while minimizing exposure to dietary toxins, potentially contributing to survival and continuing to influence modern eating behavior through circadian–sensory–nutritional interactions.

## 1 Introduction

The sense of taste acts as a gatekeeper, determining whether substances entering the mouth should be ingested or rejected (Liman et al., 2014; Yarmolinsky et al., 2009). Sweet taste signals carbohydrate-derived energy to promote food intake, whereas bitter taste helps detect toxic compounds and induces responses that halt feeding. Accordingly, taste sensitivity plays a key role in regulating dietary intake (Harnischfeger & Dando, 2021). A persistent endogenous rhythm appears to influence the taste sensitivity, thereby shaping taste perception and choice of foods across daily meals (Costanzo, 2024; Fujimura et al., 1990; Nakamura et al., 2008). Understanding oscillations in taste sensitivity may provide a perceptual basis for regulating eating behavior and optimizing dietary management across the day.

Human olfactory and auditory sensitivity reportedly increases in the evening relative to the morning (Mai et al., 2023; Marshall & Donchin, 1981) potentially conferring an advantage for detecting threats in low-light environments. By contrast, there is no consensus on diurnal variation in taste sensitivity (Costanzo, 2024), as studies of sweet taste report peak discrimination either in the morning (Nakamura et al., 2008) or superior performance at noon relative to the morning (Goetzl et al., 1950). Reports on salt sensitivity are similarly inconsistent: some indicate that it increases during the day (Irvin & Goetzl, 1952), whereas others report higher sensitivity in the evening (Fujimura et al., 1990). These inconsistencies likely reflect differences in methodologies including evaluation approaches, stimulus substances or concentration across studies, and data on bitterness, sourness, and umami remain spars (Costanzo, 2024). Circadian control of human taste function remains largely unexplored, in contrast to the well-characterized circadian regulation of feeding-related endocrine and metabolic processes (Bodosi et al., 2004; Scheer et al., 2013; Templeman et al., 2021; Trebucq et al., 2023).

A recent study has begun to uncover the circadian modulation of taste sensitivity in rodents, revealing enhanced responses to sweet, bitter and umami stimuli during the light phase, with these rhythms disappearing after knockdown of a core clock gene (Matsu-ura et al., 2025). This finding is notable because sweet, bitter, and umami transduction commonly depends on signaling pathways in type II taste cells within taste buds. However, in nocturnal mice, the enhancement of taste sensitivity during the light phase, which corresponds to their inactive or sleep period, appears to confer no obvious advantage, and its evolutionary significance remains unclear. Moreover, the limited number of comparable studies restricts progress in mechanistic and functional analyses of taste rhythms. A circadian rhythm in taste sensitivity likely exists across mammals, including humans and rodents, although its characteristics remain poorly defined. Clarifying taste function, as the gatekeeper of feeding, may help explain balanced eating behavior and support efforts to address obesity and lifestyle-related diseases.

In the present study, we characterized the overall pattern of human taste rhythms by assessing changes in taste detection, discrimination, intensity, and preference across daily meals. Additionally, testing rats during light and dark phases, we compared licking responses to solutions representing the five basic tastes. We hypothesized that if taste sensitivity has evolved adaptively for feeding behavior, the taste sensitivity exhibits opposing circadian patterns in diurnal humans and nocturnal rats, and indeed found that sweet and bitter sensitivity increased around lunchtime in humans and during the dark phase in rats. Comprehensive expression analysis of rat taste bud cells also provided multiple lines of evidence suggesting enhanced signaling in type II taste cells during the dark phase. In both humans and rats, heightened sensitivity to sweet and bitter tastes during the active phase may represent an adaptive mechanism that facilitates nutrient detection while avoiding toxins. The existence of rhythmic taste sensitivity provides insight into the long evolutionary history through which mammals established safe feeding strategies for survival.

## 2 Results

### Circadian enhancement of human taste intensity at noon

Several methods are employed to measure normal taste ability in humans (Costanzo, 2024; Fujimura et al., 1990; Goetzl et al., 1950; Irvin & Goetzl, 1952; Nakamura et al., 2008). The simple approach evaluates the subjective intensity of a taste sample placed in the mouth or assesses changes in perceived taste strength after expelling the sample (aftertaste) (Nakagawa et al., 1996). Detection and recognition thresholds are also widely applied (Irvin & Goetzl et al, 1952; Nakamura et al., 2008). The detection threshold is the minimum concentration required to perceive any taste sensation, whereas the recognition threshold represents the lowest concentration at which taste quality can be reliably identified. Most previous studies have used only one of these tests (Costanzo, 2024). Because few studies have applied multiple approaches, methodological differences likely contribute to inconsistencies among human trials. Therefore, to comprehensively capture changes in taste sensitivity, we performed above four distinct taste test types across the five basic tastes three times daily (08:00, 12:00, and 19:00; Fig. 1a).

**Fig. 1.**
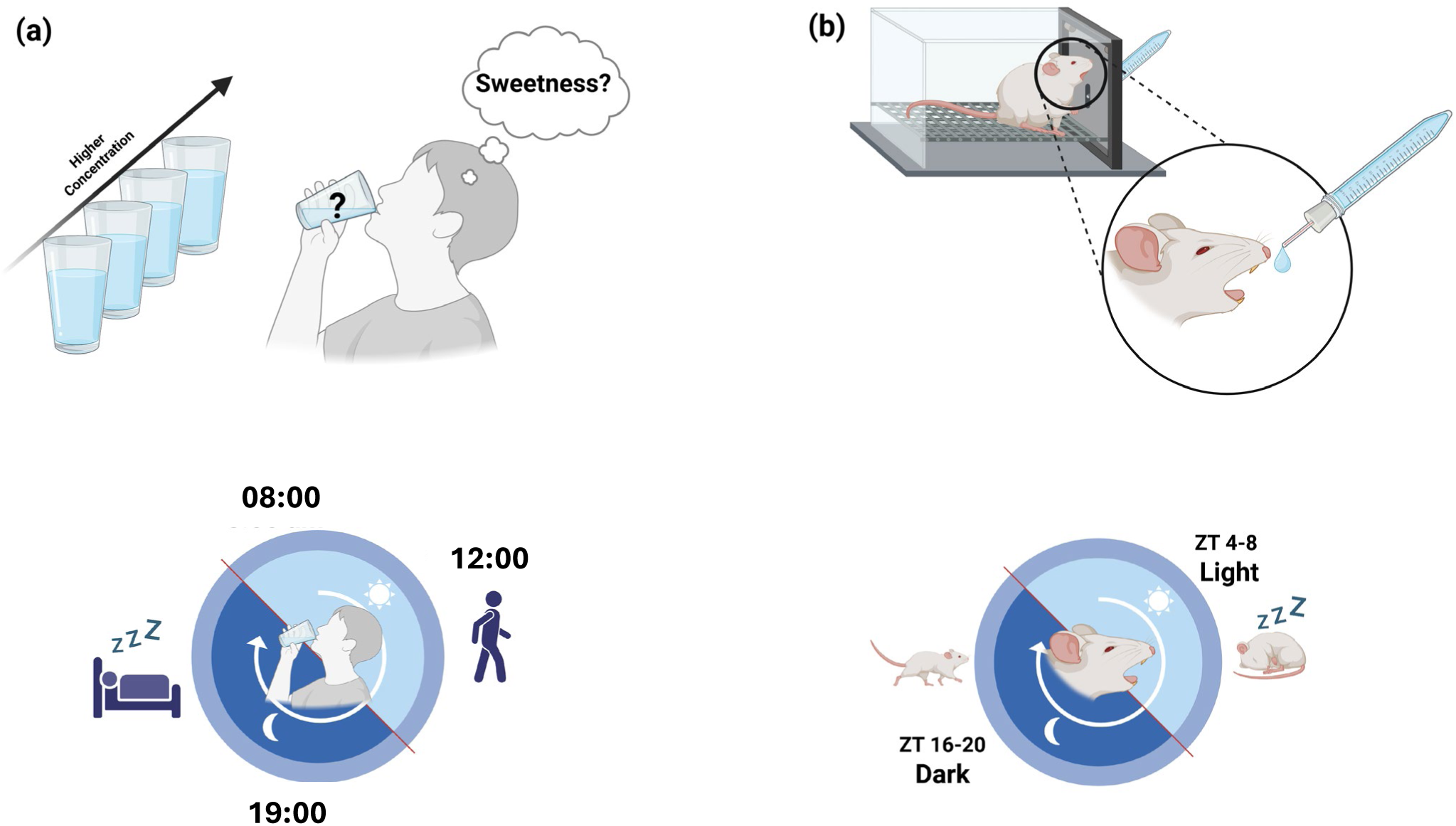
Schematic images of the taste test in humans and rats. **(a)** Human experiments used five concentration levels of aqueous solutions for the five basic tastes: granulated sugar for sweet, salt for salty, citric acid for sour, caffeine for bitter, and monosodium glutamate for umami. Participants rated taste intensity (0–5 scale), identified the taste, and selected the most preferred concentration; aftertaste intensity of the highest concentration was recorded every 10 s for 120 s. Tests were conducted at 08:00, 12:00, and 19:00. **(b)** Rat experiments used four concentrations of aqueous solutions for five tastes: sucrose for sweet, NaCl for salty, denatonium for bitter, citric acid for sour, and monosodium glutamate for umami. Citric acid and denatonium were dissolved in 250 mM sucrose to motivate licking, and licks during 10 s from first contact were measured via a sipper tube. Tests were conducted during the light (ZT4–8) and dark (ZT16–18) phases.

Circadian rhythms in taste intensity were observed for sweetness and bitterness among the five basic tastes. The intensity score for the lowest sweet concentration (0.125%) was significantly higher at noon compared with the morning or evening (p < .05; Fig. 2a). For bitterness, noon evaluation scores exceeded evening values across concentrations (p < .05; Fig. 2d). For the remaining tastes (salty, sour, and umami) scores increased only with concentration and showed no differences across testing times (Fig. 2b, c, and e, respectively). A rhythmic pattern was also evident in the aftertaste test for bitterness, where the aftertaste ratio at noon significantly exceeded that in the evening (Fig. 3d). Compared with dinner time, bitterness perception was generally stronger at lunchtime, with the sensation also persisting longer in the mouth.

**Fig. 2.**
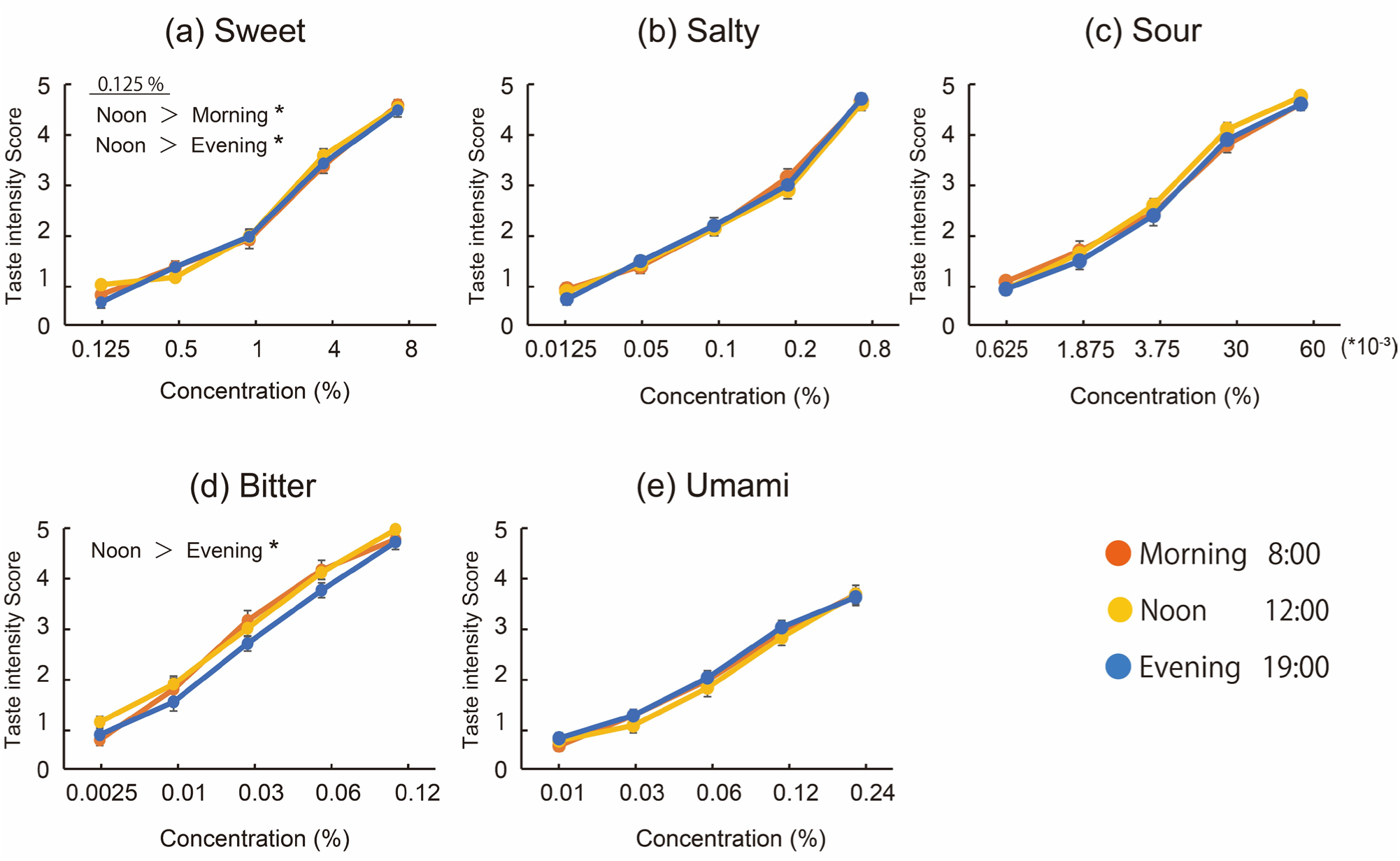
Subjective intensity scores for the five basic tastes at three daily time points in humans. Two-way ANOVA with the Shaffer post hoc test was applied (n = 19). Ratings increased with concentration across all tastes. Significant interactions and main effects were observed for sweet and bitter tastes at specific concentrations. **(a)** Sweet: concentration × time interaction trend (F_(8, 144)_ = 1.74, p < .1) indicated that at 0.125%, noon ratings were higher than morning and evening ratings (both p < .05). **(d)** Bitter: main effect of time (F_(2, 36)_ = 2.91, p < .1) showed higher scores at noon than in the evening (p < .05) regardless of concentration. **(b, c, e)** No significant time effects were observed for salty, sour, and umami solutions. Error bars represent mean ± SEM. *P < .05.

**Fig. 3.**
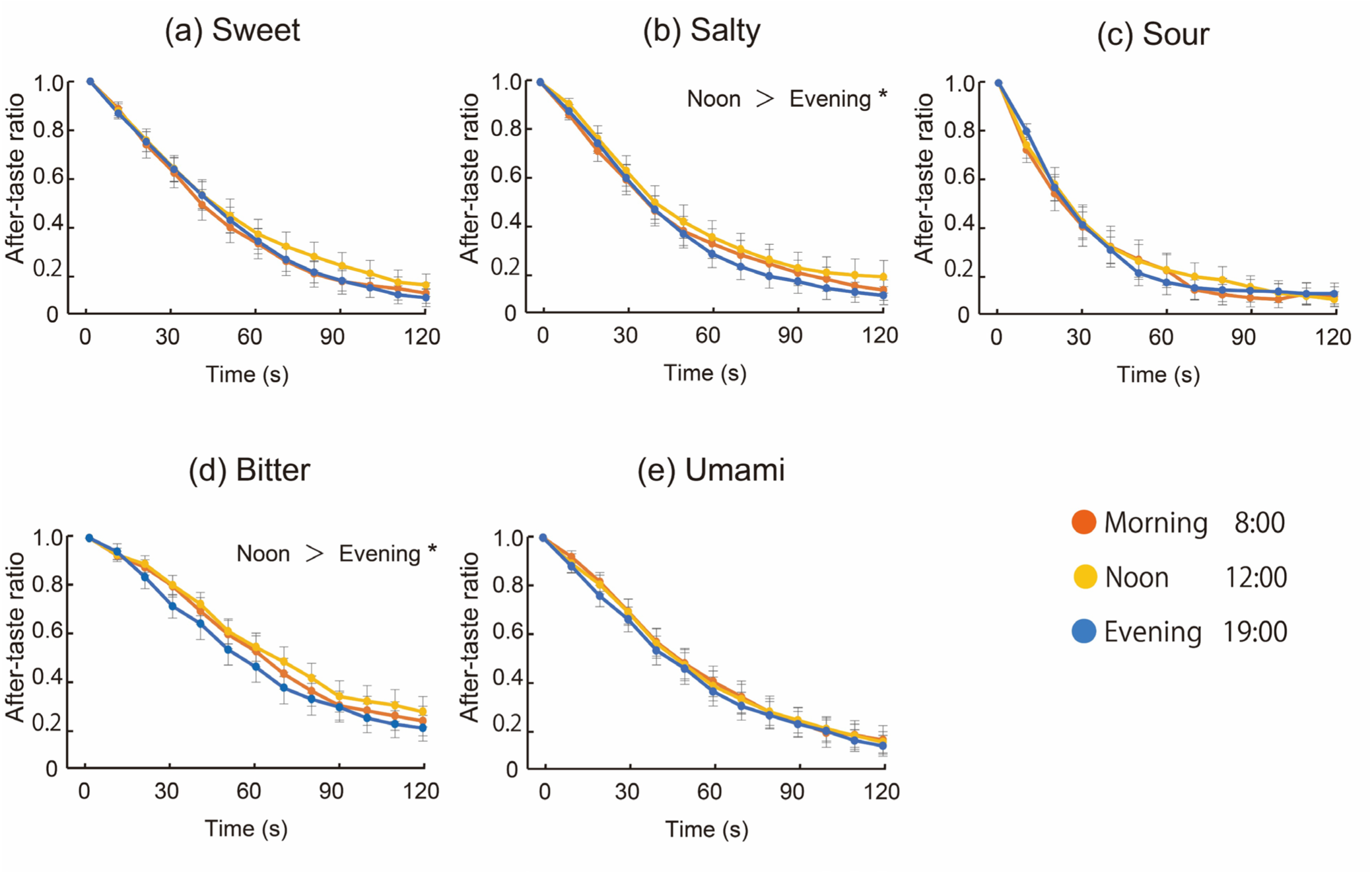
Ratios of aftertaste scores for five taste solutions at three daily time points in humans. After holding the highest-concentration solution in their mouths for 10 s, participants rated aftertaste intensity every 10 s for 120 s. Ratios were calculated relative to the initial score immediately after expectoration (0 s). Two-way ANOVA was conducted (n = 19). Ratios decreased over time for all tastes. Significant main effects of time were detected for salty (F_(2, 36)_ = 3.87, p < .05) and bitter (F_(2, 36)_ = 3.57, p < .05). **(b)** Salty: noon ratios were significantly higher than evening ratios across the 120-s rating period. **(d)** Bitter: noon ratios were significantly higher than evening ratios (p < .05). **(a, c, e)** No significant differences were observed for sweet, sour, and umami solutions. Error bars represent mean ± SEM. *P < .05.

### Greater taste intensity at lunchtime likely reflects increased sensitivity

Next, we tested whether circadian variation in taste sensitivity could be detected using threshold measurements. For sweetness and bitterness, the results were consistent with the increased intensity at noon described above. The detection threshold for sweetness ranged from 0.125% to 0.5%, with the average concentration significantly lower at noon relative to the morning or evening (p < .05; Fig. 4a, left; refer to Extended Data Fig. 1 for individual data). In contrast, the recognition threshold varied widely among individuals, ranging from 0.125% to 4%, and showed no significant daily change (Fig. 4a, middle). Preferred sweetness concentration also remained stable throughout the day (Fig. 4a, right). For bitterness, results were consistent across the three indices. The detection threshold showed a clear diurnal rhythm. Participants detected bitterness at lower concentrations and identified it more accurately at noon compared with the morning or evening (p < .05; Fig. 4d, left and middle). Accordingly, the acceptable bitterness concentration tended to drop at noon (p < .1; Fig. 4d, right). Therefore, human taste perception appears to include a mechanism that increases sensitivity during the daytime, making bitterness easier to detect. A similar mechanism may exist for sweetness but it was evident only at extremely low concentrations. Together, these tests demonstrate circadian rhythms in sensitivity to both sweet and bitter tastes. For the remaining three tastes, detection thresholds showed no significant daily variation. Meanwhile, the recognition threshold for umami was significantly higher at noon than in the morning (p < .05; Fig. 4e, middle), leading to a preference for higher umami concentrations during the day (p < .1; Fig. 4e, right).

**Fig. 4.**
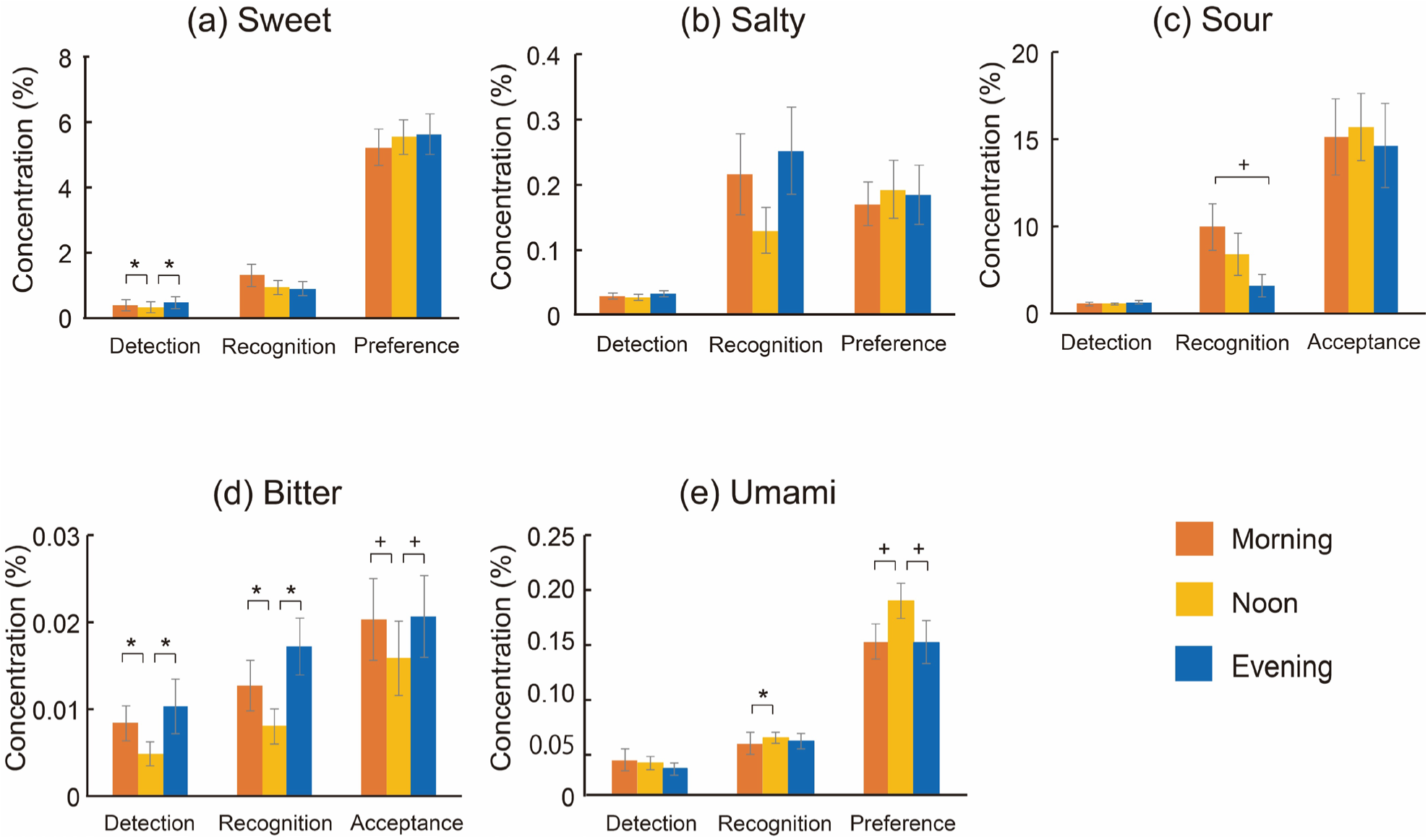
Detection threshold, recognition threshold, and preferred concentrations for five tastes at three daily time points. One-way ANOVA with the Shaffer post hoc test was applied. **(a)** Sweet: detection threshold differed significantly (F_(2, 36)_ = 5.45, p < .01), and detection concentration was significantly lower at noon compared with the morning and evening (both p < .05). **(b)** Salty: no significant differences were detected. **(c)** Sour: recognition threshold tended lower in the evening relative to the morning (F_(2, 36)_ = 2.69, p < .1). **(d)** Bitter: detection (F_(2, 36)_ = 3.33, p < .05), recognition (F_(2, 36)_ = 5.76, p < .1), and preferred (F_(2, 36)_ = 2.93, p < .10) concentrations varied; detection and recognition thresholds were significantly lower at noon than in the morning (p < .05) or evening (detection: p < .05; recognition: p < .01); preferred concentration tended lower at noon relative to the morning or evening (both p < .1). **(e)** Umami: recognition threshold (F_(2, 36)_ = 3.74, p < .05) was significantly higher at noon than in the morning (p < .05); preferred concentration (F_2, 36_ = 3.44, p < .05) tended higher at noon relative to the morning or evening (both p < .1). Recognition threshold for salty: n = 17 (two participants did not accurately recognize the taste, even at the highest concentration); all others: n = 19. Error bars represent mean ± SEM. *P < .05; ^+^p < .1.

### Independence of taste rhythms from physiological and psychological rhythms

We next examined whether physiological and psychological variables influenced taste sensitivity. Body temperature, saliva volume, subjective hunger, and fatigue levels were measured during the taste tests. Body temperature (p < .5) and subjective hunger (p < .1) increased from morning to noon, whereas fatigue scores increased from noon to evening (p < .5; Extended Data Table 1). Nevertheless, none of these parameters showed a noon-specific increase or decrease relative to morning or evening values, indicating dynamics distinct from the observed taste rhythms.

### Taste sensitivity rhythms shared across species

To determine whether taste rhythms similar to those observed in humans occur in other mammals, we conducted taste tests in rats. We compared the licking behavior of nocturnal rats toward the five basic tastes during light and dark phases by measuring the number of licks directed toward each taste solution over 10 s (Fig. 1b). When taste sensitivity is heightened, licking is expected to start to increase at lower concentrations for preferred solutions (e.g., sweet) and decrease at lower concentrations for aversive solutions (e.g., bitter).

Figure 5 shows licking rates for each solution. As in the human experiments, significant differences in response to sweet and bitter tastes were observed across the day. For bitterness, even under low-concentration conditions, licking rates during the dark phase (Zeitgeber Time: ZT16–20) were reduced compared with those during the light phase (ZT4–8) (p < .05; Fig. 5d), indicating increased bitter sensitivity and stronger aversion. The licking behavior towards bitterness was also reflected in water consumption (Fig. 6d). During the active dark phase, intake of the 0 mM bitter solution tended to exceed that during the light phase, whereas intake was strongly suppressed for the 0.5 mM solution (p < .1; Fig. 6d). These results indicate that adding 250 mM sucrose to denatonium increased licking toward the bitter solution during the dark phase (0 mM bitter solution), whereas adding 0.5 mM denatonium eliminated this sugar-driven effect and produced strong aversive behavior. Overall, licking behavior toward bitter solutions consistently indicates enhanced bitter sensitivity during the dark phase. Although sweetness sensitivity also increased during the dark phase, the rhythm was less clear. Licking rates and water consumption were elevated during the dark phase at higher concentrations (p < .05; Fig. 6a), but no significant differences appeared at lower concentrations (10 mM). Increased sensitivity to sweetness would be expected to increase licking rates at lower concentrations; however, this pattern was not observed here.

**Fig. 5.**
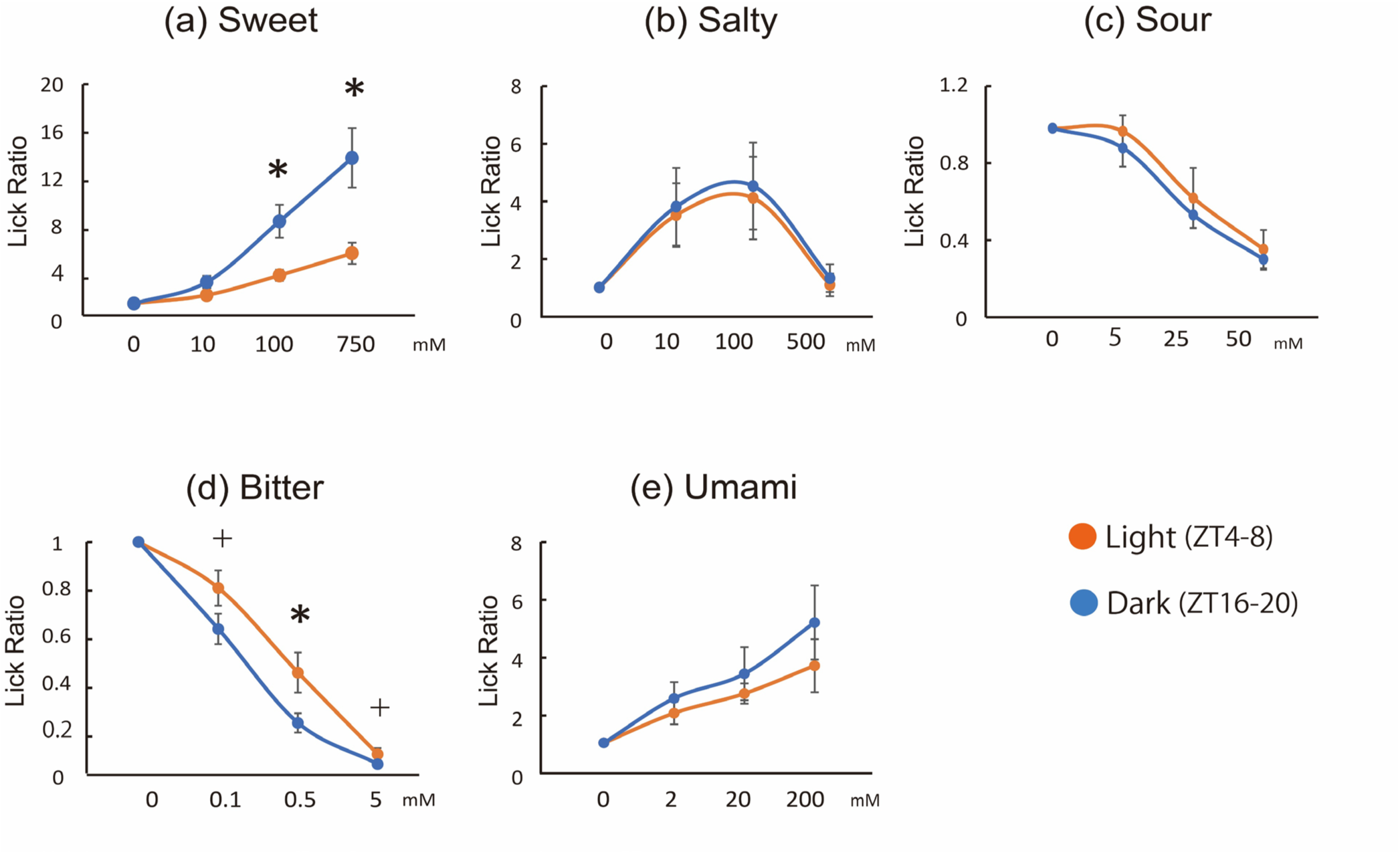
Lick ratios for five taste solutions during light and dark phases in rats. Lick ratios were calculated relative to baseline licks for the 0 mM solution. Two-way ANOVA with the Shaffer post hoc test was applied. Significant concentration × time interactions were observed for sweet (n = 11, F_(3, 30)_ = 4.47, p < .05) and bitter (n = 14, F_(3, 39)_ = 3.39, p < .05) solutions. **(a)** Sweet: dark-phase lick ratios were significantly higher than light-phase ratios at 100 and 750 mM (both p < .05). **(d)** Bitter: dark-phase ratios were lower than light-phase ratios at 0.1 mM (p < .1), 0.5 mM (p < .05), and 5 mM (p < .1). **(b, c, e)** No significant differences were observed for salty, sour, and umami solutions (n = 14 each). Error bars represent mean ± SEM. *P < .05; ^+^p < .1.

**Fig. 6.**
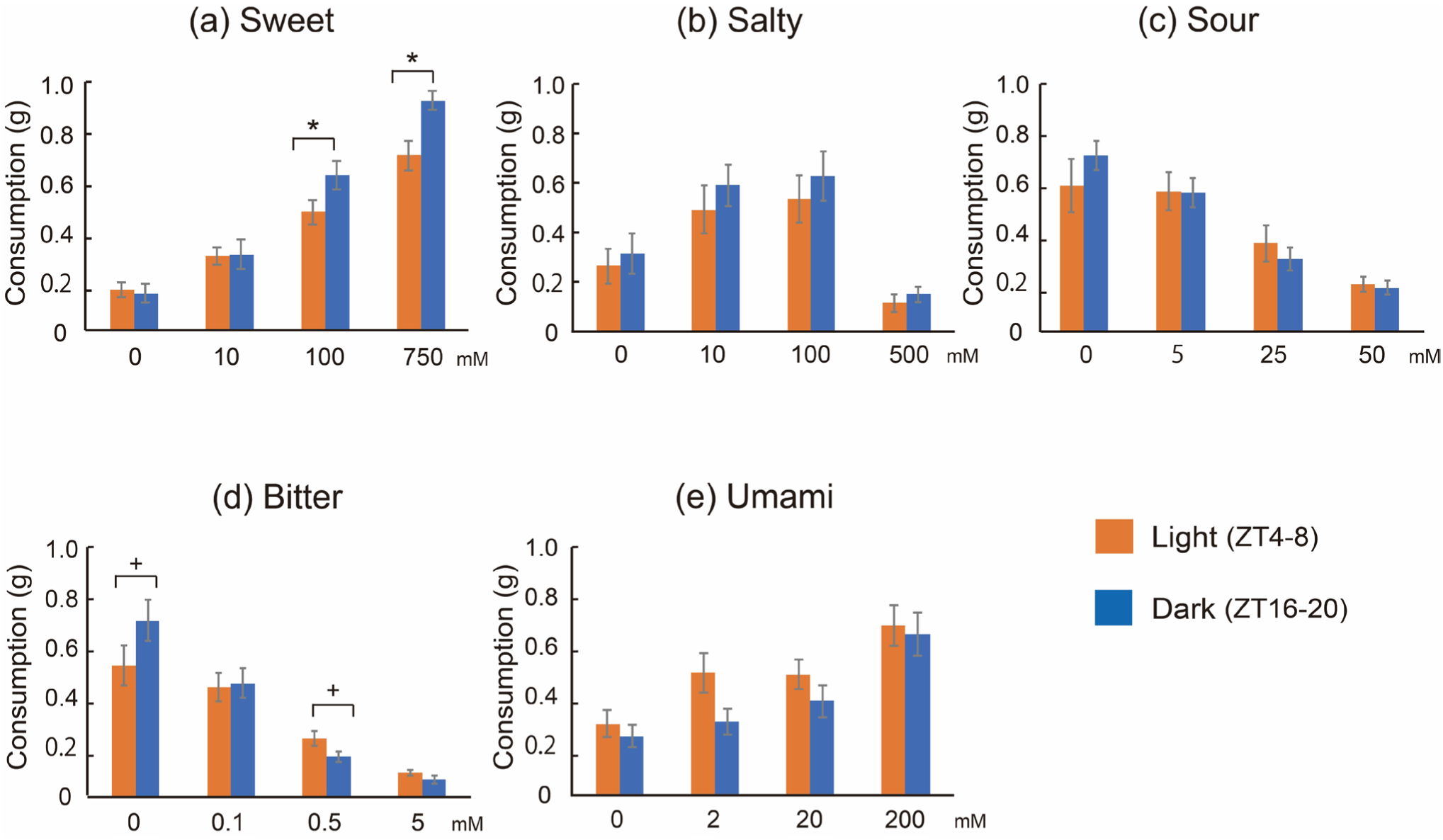
Consumption levels of the five taste solutions during light and dark phases in rats. Two-way ANOVA with the Shaffer post hoc test were applied. Significant concentration × time interactions were observed for sweet (n = 11, F_(3, 30)_ = 6.76, p < .01) and bitter (n = 14, F_(3, 39)_ = 5.04, p < .01) solutions. **(a)** Sweet: dark-phase consumption was significantly higher than light-phase consumption at 100 and 750 mM (both p < .05). **(d)** Bitter: dark-phase consumption tended higher relative to light-phase consumption at 0 mM (p < .1) but tended lower at 0.5 mM (p < .1). **(b, c, e)** No significant differences were observed for salty, sour, and umami solutions (n = 14 each). Error bars represent mean ± SEM. *P < .05; ^+^p < .1.

### Independence of blood glucose, appetite-related hormones, and taste rhythms

After completing all licking tests, blood samples were collected from rats during the light or dark phases (ZT 4–8 and ZT 16–18, respectively) to assess glucose and appetite-related hormone levels. These physiological indicators have been reported to influence feeding behavior and sweet taste sensitivity (Kawai et al., 2000; Williams & Elmquist, 2012). No significant differences were observed between phases (Extended Data Table 2). These results provide no evidence that time-dependent changes in blood components measured here explain fluctuations in taste sensitivity.

### Constant light rearing eliminates the circadian rhythm of taste

Circadian rhythms are normally synchronized to the external light–dark cycle and are substantially disrupted under conditions of continuous light (Gale et al., 2011; Ansarin et al., 2023). Therefore, we investigated whether taste sensitivity rhythms are lost under continuous light exposure. Naïve rats, which exhibited taste rhythms for sweetness and bitterness under a 12/12-h light/dark cycle, were housed under constant light for 24 h to disrupt circadian rhythms, after which licking tests were repeated. Four weeks of continuous light eliminated the dark-phase increases in food intake and total distance traveled, consistent with circadian disruption (Extended Data Fig. 2). Licking behavior also lost its light–dark differences (Fig. 7c, d). Under a 12/12-h light/dark cycle, lick rates during the dark phase were significantly higher than those during the light phase for all sweet concentrations (10–750 mM; p < .05; Fig. 7a), and increased licking at the lower concentration (10 mM) evidenced particularly enhanced sweet sensitivity during the dark phase. However, this trend disappeared under 24-h continuous light exposure, with no differences observed in licking measured at 12-h intervals (Fig. 7c, gray and black lines). Although lick rates for sweetness still increased with concentration, they plateaued at intermediate levels between the previous light and dark phases (Fig. 5a). The circadian rhythm of bitter sensitivity was similarly abolished, with the concentration-dependent decline in licking being blunted (Fig. 7d, gray and black lines), reflecting reduced sensitivity. To test whether hunger could restore sensitivity, rats under constant light were subjected to 24-h food deprivation, as fasting is known to promote licking behavior (Fu et al., 2019). However, licking rates remained unchanged (Fig. 7c, d; blue lines), indicating that fasting could not recover dulling of taste sensitivity.

**Fig. 7.**
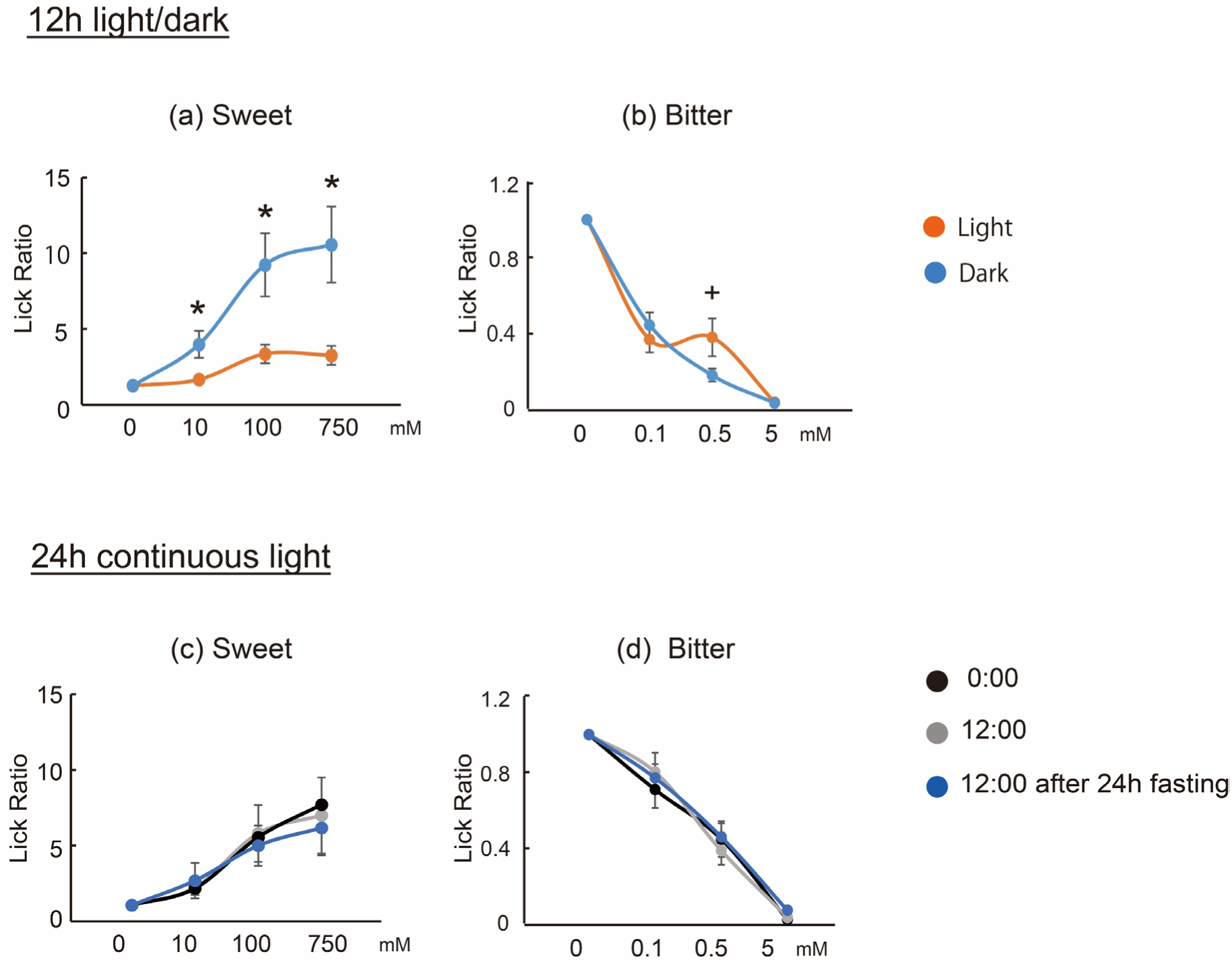
Effects of light and feeding conditions on licking behavior. Two-way ANOVA with the Shaffer post hoc test was applied. Significant concentration × time interactions were detected for sweet (n = 12, F_(3, 33)_ = 6.17, p < .01) and bitter (n = 16, F_(3, 45)_ = 3.51, p < .05) solutions under 12/12-h light/dark conditions. **(a)** Sweet: dark-phase lick ratios were significantly higher than light-phase ratios at 10, 100, and 750 mM (all p < .05). **(b)** Bitter: dark-phase lick ratios tended lower relative to light-phase ratios at 0.5 mM (p = 0.052). **(c, d)** No significant differences were identified among time points or fasting conditions (sweet: n = 12; bitter: n = 16). Error bars represent mean ± SEM. *P < .05; ^+^p < .1.

### Heightened sweet and bitter taste sensitivity during active periods in humans and rats

In diurnal humans, sensitivity to sweet and bitter tastes increased during the day, whereas in nocturnal rats, responsiveness to these tastes rose during the dark phase. These behavioral results suggest that multiple mammalian species have evolved mechanisms to enhance detection of sweet and bitter tastes during activity. Sweet and bitter stimuli are detected by type II taste cells in tongue taste buds via G protein–coupled receptors (Liman et al., 2014; Yarmolinsky et al., 2009), implicating these cells in the circadian rhythm of taste sensitivity.

### Increased expression of type II taste cell–related genes during the dark phase

To investigate the molecular basis of circadian taste sensitivity rhythms, we analyzed time-course gene expression in type Ⅱ taste cells using RNA sequencing. A prior study comparing ZT12 and ZT0 reported higher expression of *Gnat3*, a key type II taste cell gene, and the core clock gene *Bmal1*, concluding that taste sensitivity peaks during the light phase in mice (Matsu-ura et al., 2025). However, sampling only two points limits verification of a full 24-h rhythm. Our transcriptomic analysis across four time points revealed that multiple taste receptor genes were upregulated during the dark phase (ZT16 and ZT22) relative to the light phase (ZT4 and ZT10; Figs. 8–10). Expression levels of *Tas1r3* gene that forms receptors for sweet and umami tastes on the tongue, remained low during the light phase and increased upon entering the dark phase (Fig. 8c). Among 28 Tas2r bitter taste receptor genes, >50% showed significantly higher expression during the dark phase relative to the light phase; *Tas2r108*, which binds denatonium (Liman et al., 2014) used as a bitter material in our animal study, was also significantly upregulated during the dark phase (Fig. 9). Genes involved in intracellular signal transduction likewise showed increased expression during the dark phase and remained low during the light phase (Fig. 10). Multiple core clock genes, including *Bmal1* and *Clock*, were upregulated during the dark phase (Fig. 11), whereas pan taste cell marker genes (*Krt8, Krt18*) showed stable expression levels (Extended Data Fig. 3) over 24 h in type Ⅱ taste cells. These results suggest that, in type II taste cells, gene expression of taste-related genes were upregulated during the active dark phase, thereby promoting efficient tastant capture and signal transmission to the brain. Our molecular analyses are consistent with behavioral evidence indicating that sensitivity to sweet and bitter tastes is heightened during the mid-dark phase (ZT16–20) compared with the light phase (ZT4–8).

**Fig. 8.**
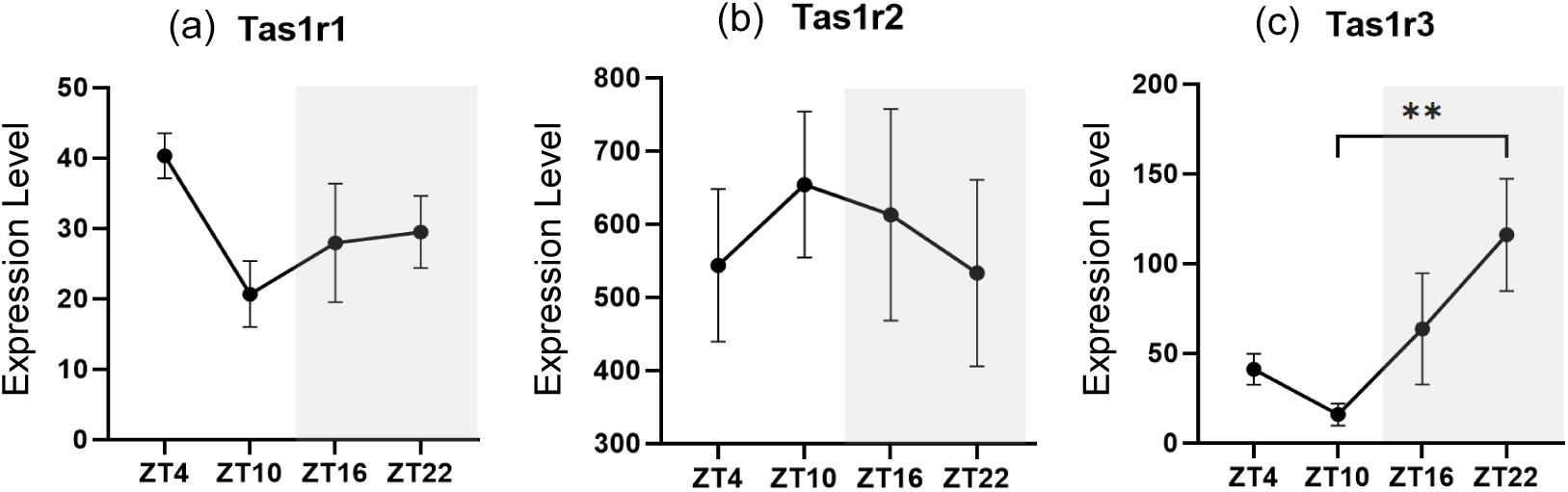
Daily fluctuations in *Tas1r* family gene expression levels in rat tongue type Ⅱ taste cells. Kruskal–Wallis test with Dunn’s multiple comparison was applied (n = 6 per condition). Error bars represent mean ± SEM. **P < .01.

**Fig. 9.**
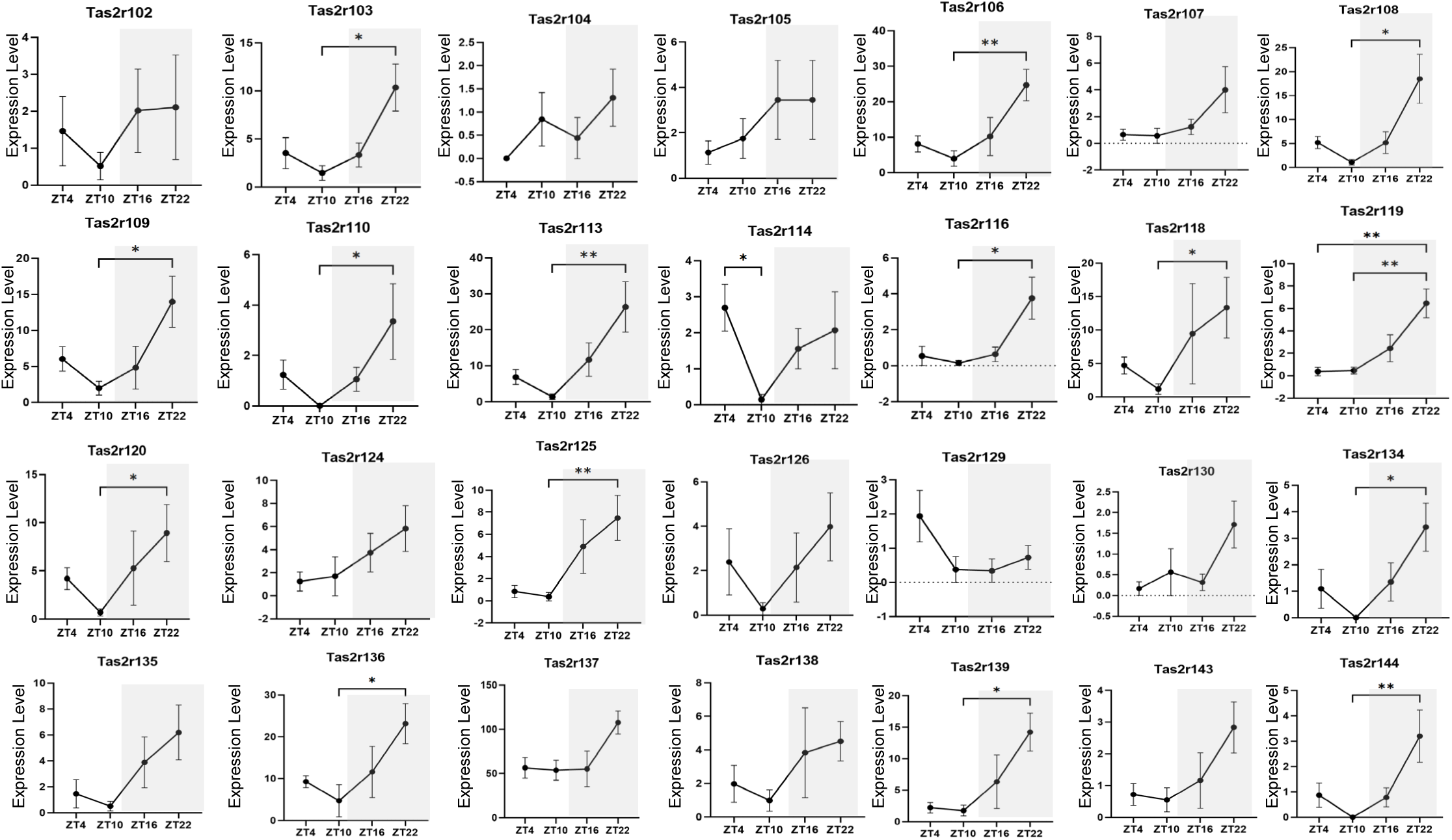
Daily fluctuations in *Tas2r* family genes expression levels in type II taste cells. Kruskal-Wallis test with Dunn’s multiple comparison was applied (n = 6 per condition). Error bars represent mean ± SEM. **P < .01; :p < .05.

**Fig. 10.**
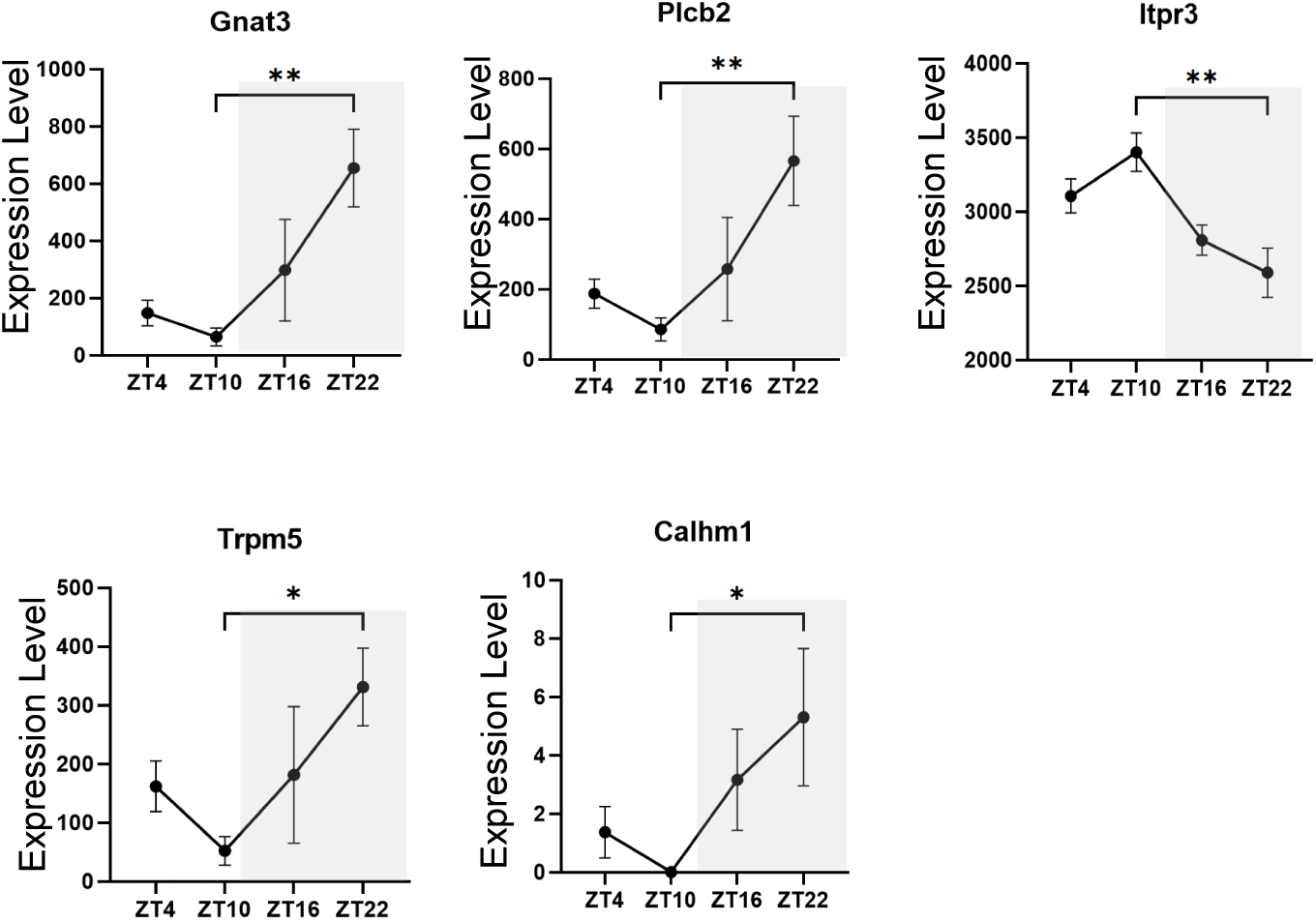
Daily fluctuations in the expression levels of genes involved in intracellular signal transduction in type Ⅱ taste cells. Kruskal–Wallis test with Dunn’s multiple comparison was applied (n = 6 per condition). Error bars represent mean ± SEM. **P < .01; *p < .05.

**Fig. 11.**
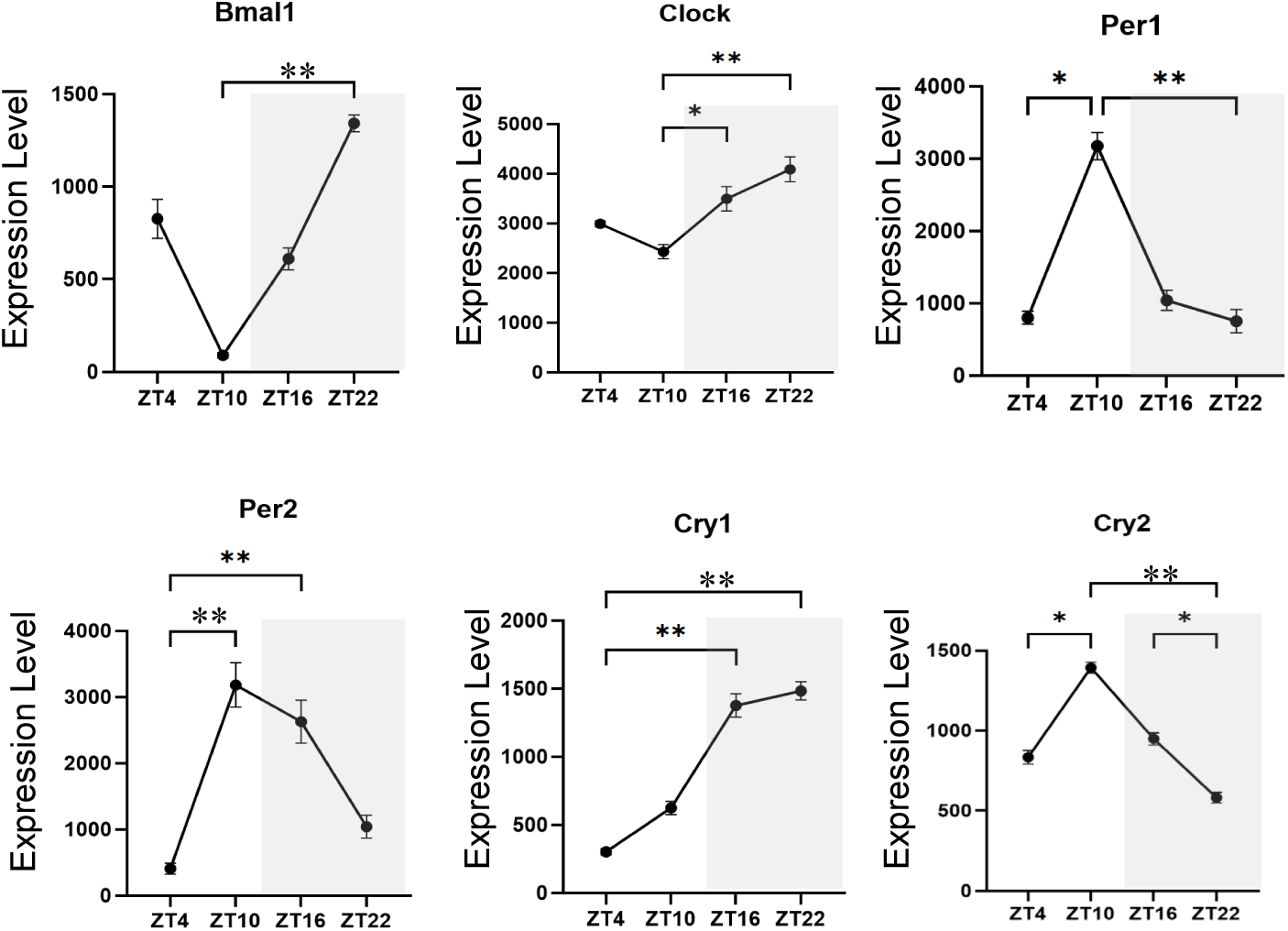
Daily fluctuations in clock gene expression levels. Kruskal–Wallis test with Dunn’s multiple comparison was applied (n = 6 per condition). Error bars represent mean ± SEM. **P < .01; *p < .05.

## 3 Discussion

Our study provides consistent evidence that taste sensitivity is enhanced during active periods in both humans and rats, potentially mediated by circadian regulation of taste-related gene expression in tongue taste buds. In diurnal humans, the intensity of sweetness and bitterness is perceived more strongly, and detection or recognition sensitivity is heightened at lunchtime compared to morning or evening. In nocturnal rats, behavioral changes indicative of increased taste sensitivity were observed during the dark phase, manifested as an enhanced preference for sweet solutions and more pronounced avoidance of bitter solutions. Furthermore, consistent with behavioral changes, the expression levels of receptors and intracellular signaling genes involved in sweet and bitter taste reception were significantly higher during the dark phase compared with the light phase in taste buds. Such circadian rhythms may be advantageous in enabling efficient energy acquisition while limiting exposure to harmful substances. Moreover, this taste rhythm may be shared across diel activity patterns, implicating a role in enhancing mammalian survival over evolutionary history.

Notably, convergent evidence across species was observed for rhythmic sensitivity to the shared taste modalities of sweetness and bitterness. However, this circadian pattern differs from that reported for auditory and olfactory systems, which are more sensitive in the evening than in the morning in humans (Mai et al., 2023; Marshall & Donchin, 1981) and rodents (Granados-Fuentes et al., 2011: Halberg et al., 1958; Takeuchi et al., 2023). Circadian regulation with olfactory sensitivity likely evolved to detect threats occurring in darkness (Granados-Fuentes et al., 2011; Mai et al., 2023), whereas gustatory sensitivity seems to be adapted to optimize foraging efficiency and safety, peaking during each species’ active period. These findings indicate that circadian modulation of sensory functions might be neither uniform nor universal, but has evolved with functional specificity.

The circadian rhythm of taste sensitivity appears to be predominantly mediated by peripheral taste receptor cells in the taste buds, potentially distinguishing its regulation from olfactory processing (Granados-Fuentes et al., 2011; Takeuchi et al., 2023). In olfaction, involvement of central mechanisms has been suggested, including modulation at the level of the olfactory bulb and piriform cortex (Granados-Fuentes et al., 2011; Takeuchi et al., 2023). Consequently, odor stimuli are not expected to be differentially regulated at the receptor level, and circadian fluctuations likely affect olfactory signals more uniformly as they reach the central nervous system (Takeuchi et al., 2023). In contrast, circadian regulation of taste was not uniform across the five basic taste modalities, indicating selective regulation within peripheral taste cells. Saltiness and sourness did not show clear daily modulation was detected in our experiments. These differences among the five tastes are consistent with prior mouse research (Matsuura et al., 2025). Sensitivity to saltiness and sourness can change with exercise and salt intake (Gauthier et al., 2020; Brot et al., 2000). Because sodium and hydrogen ions contribute directly to physiological homeostasis, sensitivity to saltiness and sourness may be regulated in a more rapid and flexible manner in response to ion and water balance, with potentially less dependence on circadian control. Such flexible modulation for each taste modality suggests that peripheral regulation may be favored to control sensitivity.

Our human threshold measurements, together with gene expression analyses, further suggest that circadian modulation of taste sensitivity arises primarily at the peripheral level. Circadian rhythmicity was observed in sweet taste detection thresholds and intensity ratings, but not in recognition thresholds or preferred concentrations. As basic sensory functions, detection thresholds and intensity ratings more directly reflect peripheral gustatory input and may therefore be particularly sensitive to circadian modulation, relying on relatively limited higher-order processing within the primary gustatory cortex, including the insular and anterior cingulate cortices. In contrast, evaluating taste quality and preference requires processing across broader brain regions, including the orbitofrontal cortex and anterior cingulate cortex (Avery et al., 2020; Rolls et al., 2003). Increasing complexity of central processing can diminish differences in peripheral signal strength, thereby obscuring circadian rhythms in taste perception. Furthermore, the cognitive threshold concentration was 3.4-fold greater than the detection threshold, and the preferred concentration was ∼20-fold greater. These values suggest that humans are routinely exposed to sweet concentrations far exceeding the detection threshold in everyday diets. The concentration at which sweetness is recognized, as well as the preferred level, may also be influenced by dietary habits (Costanzo, 2024; Harnischfeger & Dando, 2021). These observations suggest that circadian variation in sweetness sensitivity plays only a modest role in shaping dietary behavior in modern humans, whose diets have become increasingly rich in sweetness through dietary modernization (Rao, 2026).

Compared with sweetness sensitivity, bitter taste perception appears more closely linked to intake decisions even in modern diet. The concentration at the bitterness perception threshold was 1.6-fold greater than the detection threshold, and the acceptable concentration was 2.4-fold greater, the differences between these concentrations were relatively small. Differences in peripheral signal intensity are reflected in detection thresholds and can lead to fluctuations in the acceptable level of bitterness. Because the ability to perceive bitterness is closely tied to toxin avoidance, the strong link between peripheral detection sensitivity and central feeding decisions might have been evolutionarily preserved. In our human experiment, sensitivity differences were consistently greater between noon and evening than between noon and morning, likely owing to the longer interval separating the noon and evening tests. Increasing the number of measurements per day beyond three would enable more precise identification of the peak taste sensitivity period.

Although no clear circadian rhythm was detected in umami sensitivity, our molecular data suggest that umami—mediated by type II taste cells—may still be enhanced during the active period. This possibility warrants reevaluation using a broader concentration range and/or alternative umami stimuli. Beyond monosodium glutamate (used here), several umami-active compounds, including inosinate and guanylate, commonly co-occur in foods, and their combination can synergistically increase perceived umami intensity (Araujo et al., 2003; Li et al., 2002). Accordingly, future studies employing mixtures of umami compounds could help determine whether circadian timing modulates the synergistic perception of umami. Interestingly, our human experiment showed that daytime sensitivity decreased for the umami recognition threshold and preferred concentration, rather than for umami detection. This pattern could be related to higher cognitive processing in the central nervous system rather than to alterations in peripheral sensitivity. However, research on umami sensitivity remains limited (Costanzo, 2024), and the cognitive mechanisms underlying this phenomenon are not yet explained.

In human experiments overall, circadian rhythmicity was more clearly detected in basic perceptual judgments of sweetness and bitterness, consistent with molecular evidence indicating predominantly peripheral regulation within taste cells. Previous studies have often relied on cognitive thresholds (Nakamura et al., 2008; Goetzl et al., 1950; Irvin & Goetzl, 1952; Fujimura et al., 1990), which depend on central processing and may contribute to inconsistent results. By systematically conducting multiple tests, our study helps reconcile these discrepancies and shows that sweet and bitter sensitivity peaks during the daytime in humans.

Regarding rat licking behavior, stronger preference or aversion was observed during the dark phase relative to the light phase with low-concentration sweet and bitter tastants, indicating heightened sensitivity. These behavioral changes were accompanied by elevated expression of taste-related genes in type II taste cells during the dark phase, suggesting a molecular mechanism for circadian regulation of taste sensitivity. These findings partially align with the study by Matsuura et al. (2025) on mice, which implicated that type II taste cells are involved in the circadian rhythm of taste sensitivity, although their results suggested the opposite pattern, i.e., that taste sensitivity increased during the light phase. This discrepancy may reflect differences in experimental timing. In our study, gene expression was measured at four time points (ZT4, ZT10, ZT16, and ZT22), revealing a clear rhythm in which taste-related gene transcription increased during the dark phase and remained relatively low during the light phase. In contrast, Matsu-ura and colleagues (2025) compared only two time points (ZT0 and ZT12), which may have obscured the rising trend occurring from ZT12 to ZT24 (dark period).

We found that several type II taste-cell signaling genes (*Gnat3*, *Trpm5*, and *Calhm1*) increased during the dark phase relative to the light phase, consistent with enhanced downstream signaling of sweet and bitter transduction that may contribute to the heightened sensitivity observed at this time. The dark-phase upregulation of genes such as *Tas1r3*, *Tas2r108*, and *Tas2r134* could be associated with the maintenance or renewal of taste receptor function; however, direct evidence is limited, highlighting the need for future studies. Genes involved in sour taste reception (*Pkd2l1* and *Otop1*) were upregulated during the dark phase, but the relationship between this phenomenon and circadian changes in gustatory sensitivity is unknown. It is possible that subtle circadian rhythms are embedded in the dynamic regulation of sensitivity driven by fluctuations in ion balance. Additionally, clock genes in type II taste cells also showed similar circadian oscillations, suggesting a potential link between the circadian clock machinery and taste-related gene regulation.

Continuous light exposure disrupted circadian timing in rats and abolished the circadian rhythm of taste sensitivity. These results suggest several auxiliary mechanisms linking light cycles to taste sensitivity, including modulation via central circadian control originating from the suprachiasmatic nucleus, as well as indirect modulation through behaviorally mediated feedback, such as changes in feeding timing. Elucidating the mechanism by which the light cycle influences circadian rhythms in taste sensitivity is an important challenge for future research.

The circadian rhythm of taste sensitivity did not synchronize with the fluctuations in subjective feeling of hunger in humans. Twenty-four-hour fasting in rats appeared to have little effect on licking rates. Our results were unable to demonstrate a daily rhythm in ghrelin and leptin concentrations levels in blood due to the limitations of two-point measurements (ZT4–8 and ZT16–20) and individual differences in circadian phase and feeding schedules under ad libitum feeding (Maffei et al., 1995; Tolle et al., 2002), but they also did not strongly support a connection between the circadian regulation of appetite-related hormones and taste sensitivity. These findings suggest that the observed rhythm of taste sensitivity arises largely independently of appetite-related factors. In contrast, increased licking by rats during the dark phase occurred not only at low-concentration sweet solutions but also at higher concentrations. This pattern is unlikely to reflect changes in taste sensitivity alone, and may instead indicate enhanced motivational states during the dark phase, encompassing drinking behavior associated with hunger and thirst as well as increased motivation for carbohydrate consumption (Bainier et al., 2017). Some evidences also suggest that taste sensitivity itself may be influenced by hypothalamic regulation of appetite (Fu et al., 2019; Kawai et al., 2000; Zizzari et al., 2011; Zverev, 2004). A comprehensive understanding of how taste sensitivity is translated into eating behavior requires integrated analyses across peripheral tastant-sensing pathways and central appetite-regulating neural circuits.

In conclusion, despite their contrasting diurnal and nocturnal activity patterns, humans and rodents share a common feature: sensitivity to sweet and bitter tastes increases during their respective active periods. This circadian modulation appears to be supported by regulation of taste-related gene expression in type II taste cells in the tongue. Such rhythmic variation may have evolved to help organisms efficiently locate energy sources while avoiding toxins. In modern human dietary environments, sweetness is often encountered at concentrations far exceeding detection thresholds, which might mask circadian changes in sweet taste sensitivity. In contrast, fluctuations in bitter taste sensitivity remain more perceptible and may retain greater relevance to contemporary dietary behavior. These findings suggest that circadian regulation of taste sensitivity reflects conserved molecular mechanisms across mammals while also influencing food preferences and dietary choices, highlighting an underexplored intersection involving circadian biology, sensory perception, and nutrition. Strategic leverage of taste functions that operate consciously/unconsciously across daily eating occasions may ultimately enable the development of dietary strategies that align with taste preferences while ensuring appropriate intake of essential nutrients.

## 4 Materials and Methods

### Human Experiments

#### Participants

Twenty healthy Japanese-speaking adults (8 males and 12 females; mean age, 35.7 ± 11.3 years; range, 21–57 years) were recruited in this study. Inclusion criteria were age ≥20 years and native proficiency in Japanese, while exclusion criteria included dietary restrictions due to illness, pregnancy, and a known allergy to caffeine. Taste function was assessed using the Screening Questionnaire for Loss of Taste (Malaty et al., 2013), and data from 19 participants with no evidence of taste loss were included in the analyses.

#### Taste solutions

Aqueous solutions representing five basic tastes were prepared: granulated sugar (0.125%–8% w/w) for sweet, sodium chloride (NaCl; 0.0125%–0.8% w/w) for salty, citric acid (0.000625%–0.06% w/w) for sour, caffeine (0.0025%–0.12% w/w) for bitter, and monosodium glutamate (0.01%–0.24% w/w) for umami. Each tastant was diluted with distilled water (DW), which was also used for mouth rinsing after each sample. Each solution (30 g) was sealed in a coded food-grade plastic bag and stored frozen. Before testing, samples were brought to room temperature and transferred to plastic cups for use.

#### Taste perception test schedule

Five types of taste perception tests were conducted in the following order: aftertaste, subjective intensity, detection threshold, recognition threshold, and preference (acceptability), each performed three times daily (08:00, 12:00, and 19:00). In all tests, the five solutions were presented in a predetermined random order. Participants were instructed to abstain from food and beverages (including coffee or alcohol) that could affect their physiological state for 2 h before testing. After receiving protocol instructions, participants practiced the full set of tests at least twice [once at the National Agriculture and Food Research Organization (NARO) laboratory and once at home] to ensure proficiency. Those confirmed to be proficient completed the tests at home using three sets of prepared samples at the specified times. Testing was conducted at home owing to the COVID-19 pandemic, during which social distancing was recommended. All tests were performed inside a white portable booth (50 × 70 × 60 cm) placed in a room maintained at a mean temperature of 21.6°C ± 1.7°C (± standard deviation). Female participants were tested 3–16 days after the end of the menstrual phase (Nakamura et al., 2008).

#### Amount of aftertaste

Participants held 15 mL of the highest-concentration sample for each taste in their mouths for 10 s, after which it was expectorated and aftertaste intensity evaluated for 120 s at 10-s intervals. Intensity was rated on a 12-cm line scale ranging from no taste to extremely strong taste. Each 15-mL sample was provided in a small plastic cup, and participants identified the taste before rating. Aftertaste reduction ratios were calculated relative to the score immediately after expectoration (0 s).

#### Subjective taste intensity, detection, and recognition thresholds

Each 30-mL sample for all five tastes was placed in a plastic cup. Participants tasted each solution starting from the lowest concentration and rated intensity on a scale from 0 to 5 (0: no taste; 1: very weak; 2: weak; 3: medium; 4: strong; 5: very strong taste). When a taste was detected (score ≥1), participants identified it as sweet, salty, sour, bitter, or umami. After each evaluation, the next higher concentration was tested, and the same procedure was repeated up to the highest concentration. The lowest concentration with a score ≥1 was defined as the detection threshold, and the concentration at which the taste was first correctly identified was defined as the recognition threshold.

#### Preference (acceptability) for taste

After completing threshold tests with the highest concentration samples, participants selected the most preferred (or acceptable) concentration among the five levels for each taste.

#### Body temperature, saliva volume measurement, and other questionnaires

Participants measured axillary body temperature during the taste test. Saliva volume was also assessed by chewing cotton wool (Salimetrics Children’s Swab) for 1 min. Subjective hunger and fullness were rated on a 12-cm line scale, and fatigue was evaluated using the Profile of Mood States scale short form (Yokoyama, 2006).

### Rat Experiments

#### Animal subjects

All animal procedures were approved by the University of Tsukuba Committee on Animal Research. Sexually naïve male Wistar–Imamichi rats (8 weeks old) were obtained from the Institute for Animal Reproduction (Ibaraki, Japan). Animals were housed in transparent polypropylene cages (23 × 38 × 20 cm) containing wood chip bedding under controlled temperature (23°C ± 1°C) and humidity (50% ± 10%) conditions, maintained under a 12/12-h light/dark cycle (lights on at 08:00; 100 lux during the light phase), with *ad libitum* access to food and water.

#### Licking behavior

All experiments were performed in an opaque polypropylene chamber (29 × 18 × 14 cm) with a square opening (2.1 × 1.1 cm) on the front panel that allowed the rat to extend its tongue to a stainless-steel sipper tube (44-mm long) connected to a 15-mL plastic tube containing the taste solution. Taste stimuli included sucrose (0–750 mM) for sweet, NaCl (0–500 mM) for salty, denatonium (0–5 mM) for bitter, citric acid (0–50 mM) for sour, and monosodium glutamate (0–200 mM) for umami. Citric acid and denatonium were dissolved in 250 mM sucrose solution to promote licking behavior. All solutions were presented at room temperature. Licks were recorded using a custom-built lickometer (F. A. Systems, Ibaraki, Japan) in which an optical sensor detected interruptions caused by the rat’s tongue. Before testing, rats underwent 30-min training sessions per day for 4–5 days to establish stable licking behavior, during which water and chow were restricted. On day 1, rats explored the chamber for 30 min for habituation. On day 2 and thereafter, three of six solutions (DW and five tastants) were presented in random order for 10 min. Trained rats were deprived of water for 4–6 h before testing to increase motivation for solution access.

The licking test comprised five sessions during the light phase and five during the dark phase. Each session was conducted during the light (ZT4–8) or dark (ZT16–18) phase, with one of the five tastants presented per session. The order of stimuli presentation was counterbalanced. Licks were counted for 10 s from the first contact with the sipper tube, and this measurement was repeated four times across a series of four concentrations for each tastant. Each concentration was presented twice in ascending and descending order. After testing, rats were returned to their home cages and given *ad libitum* access to water 2-h later. Lick ratios were calculated relative to baseline licking for DW (salty, sweet, and umami) or 250 mM sucrose solution (sour and bitter) during each 10-s test session. Total solution intake at each concentration was also measured. Rats that consumed large volumes of solution (including DW), regardless of concentration, due to excessive thirst were retested after 2 min of free access to DW in the home cage.

#### Circadian rhythm disruption procedure and travel distance analysis

Baseline licking responses to sweet and bitter tastants were first measured in naïve rats maintained under a standard 12/12-h light/dark cycle. To disrupt circadian rhythm, rats were then transferred at 21 weeks of age to constant illumination (80–100 lux; 24/0-h light/dark). Before and after 4 weeks of continuous light exposure, locomotor activity was recorded for 24 h using a Tapo C210 V3 Wi-Fi camera (TP-Link Systems Inc., CA, USA) to calculate travel distance. Videos were recorded at 30 fps and downsampled to 1 fps using Simple Behavioral Analysis (v2.3.2). Body part tracking was performed using DeepLabCut (v2.3.9) with a ResNet-50 network trained for 100,000 iterations on 320 labeled frames with eight landmarks (left ear, right ear, nose, mid-body, left body, right body, tail base, and tail tip). Training and test errors were 2.83 and 8.08 pixels, respectively. Total travel distance was calculated using a custom Python (v3.8.19) script. Food intake was also measured every 12 h once per week during both the 4-week continuous light period and the preceding 4 weeks. After confirming circadian disruption based on locomotor activity and feeding rhythms, licking tests were repeated to assess the effects of constant light exposure on taste sensitivity. Additionally, the licking behavior of rats with disrupted circadian rhythms was retested after 24 h of food deprivation.

#### Cardiac blood sampling, blood glucose measurement, and enzyme-linked immunosorbent assay

All procedures were performed under 5% isoflurane anesthesia after confirming loss of the righting reflex. Cardiac blood was collected using 21-gauge needles attached to 5-mL syringes during the light (ZT4–8) and dark (ZT16–18) phases. For blood glucose measurement, 200 µL of blood was immediately applied to a blood glucose meter (MEDISAFE FIT Smile Blood Glucose Meter, Terumo, Japan), and values were recorded. To measure leptin, 1 mL of blood was placed in a microcentrifuge tube and centrifuged at 3,500 rpm for 10 min at 4°C, and the serum was collected. For acylated and unacylated ghrelin assays, 2 mL of blood was collected into an EDTA-2Na tube, gently mixed, transferred to a PHMH tube, and centrifuged at 3,500 rpm and 4°C for 10 min, after which the plasma was collected. All samples were stored at −80°C until enzyme-linked immunosorbent assay (ELISA) analysis. The Mouse/Rat Leptin ELISA Kit (M1305, Morinaga, Japan), Ghrelin Easy Sampling ELISA/Mouse/Rat/Acylated Ghrelin (A05317, Bertin Bioreagent), and Ghrelin Easy Sampling ELISA/Mouse/Rat/Unacylated Ghrelin (A05318, Morinaga, Japan) were used according to the manufacturers’ instructions to determine soluble leptin and ghrelin levels.

#### Taste bud cell circumvallate papillae sampling for RNA sequencing

Following 5% isoflurane anesthesia, animals were decapitated, and circumvallate papillae were excised to isolate taste buds. Samples were immediately placed in RNAlater (Thermo Fisher Scientific, USA) and stored at −80°C for subsequent RNA sequencing.

#### RNA-sequencing analysis

Samples from each group were processed for RNA sequencing at Macrogen Japan (Tokyo, Japan) using the NovaSeq X platform. TruSeq Stranded mRNA libraries were constructed for paired-end 151-bp sequencing. Sequence quality was assessed using FastQC (v0.11.7), and transcript assembly and expression quantification were performed using StringTie (v2.1.3b). For gene expression analyses, normalized read counts were obtained using DESeq2 (v1.38.3) in R (v4.5.2).

### Statistical Analysis

Results are presented as means ± standard errors of means in each figure. Behavioral and physiological data were analyzed using one-way or two-way analysis of variance (ANOVA), with Shaffer’s method applied for multiple post hoc comparisons. Differences were considered significant at *p* ≤ 0.05, and trends were noted at *p <* 0.1. Individual gene expression differences were analyzed using the Kruskal–Wallis test followed by Dunn’s multiple comparisons test.

## Ethics Statement

The human experiment protocol was approved by the Institutional Human Research Review Board of NARO. All participants provided written informed consent prior to participation and could withdraw at any time. Participants received financial compensation for participation. All animal procedures were approved by the University of Tsukuba Committee on Animal Research and conducted in accordance with the Guidelines for the Care and Use of Laboratory Animals of the Japanese Ministry of the Environment (notification no.: 88, 2006) and related regulations. Study reporting followed the ARRIVE guidelines.

---

**Extended Data Fig. 1.**
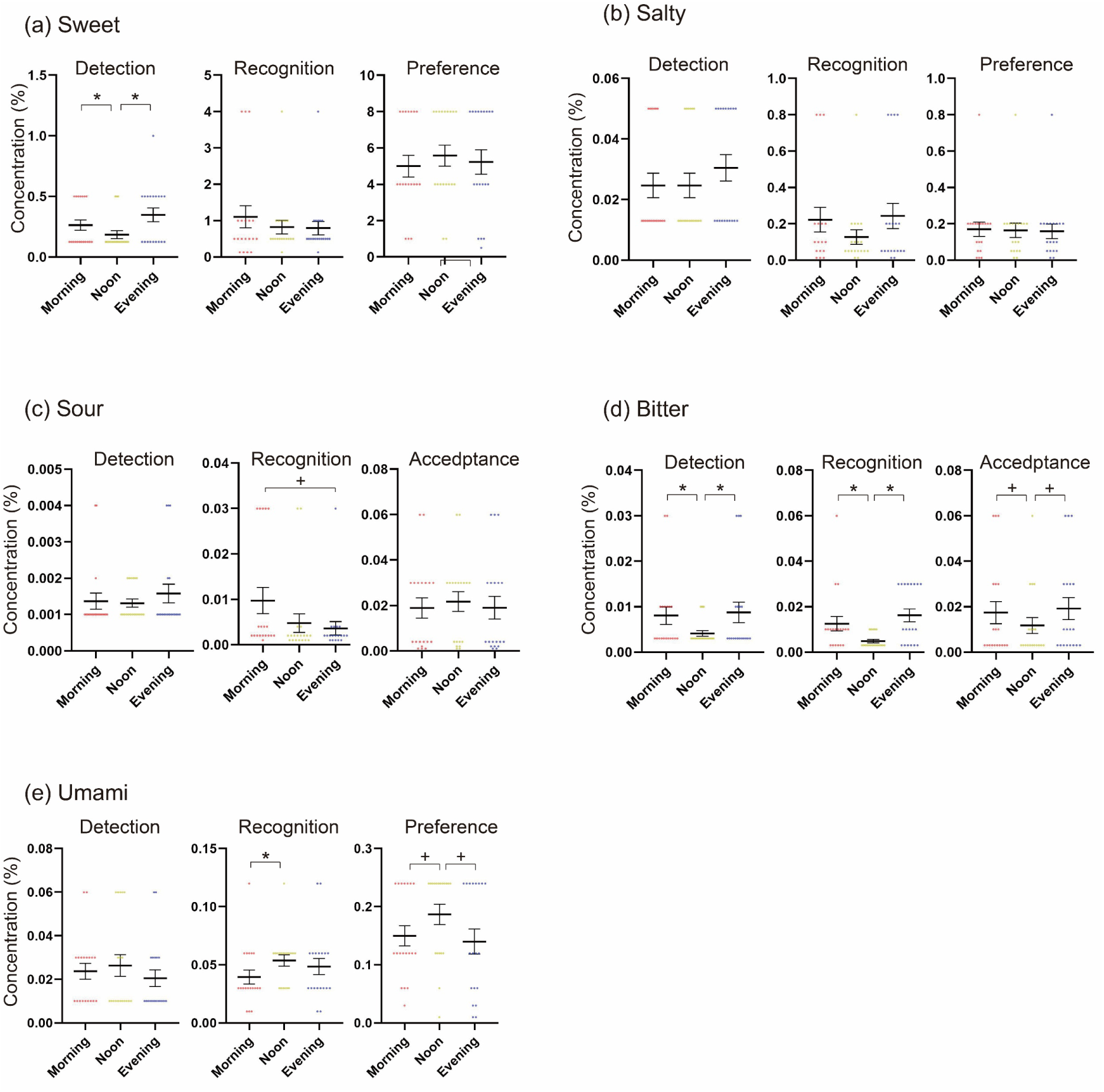
Individual differences in detection thresholds, recognition thresholds, and preferred (accepted) concentrations. Statistical results and additional details are shown in Fig. 4. Error bars represent mean ± SEM. *P < .05; ^+^p < .1.

**Extended Data Fig. 2.**
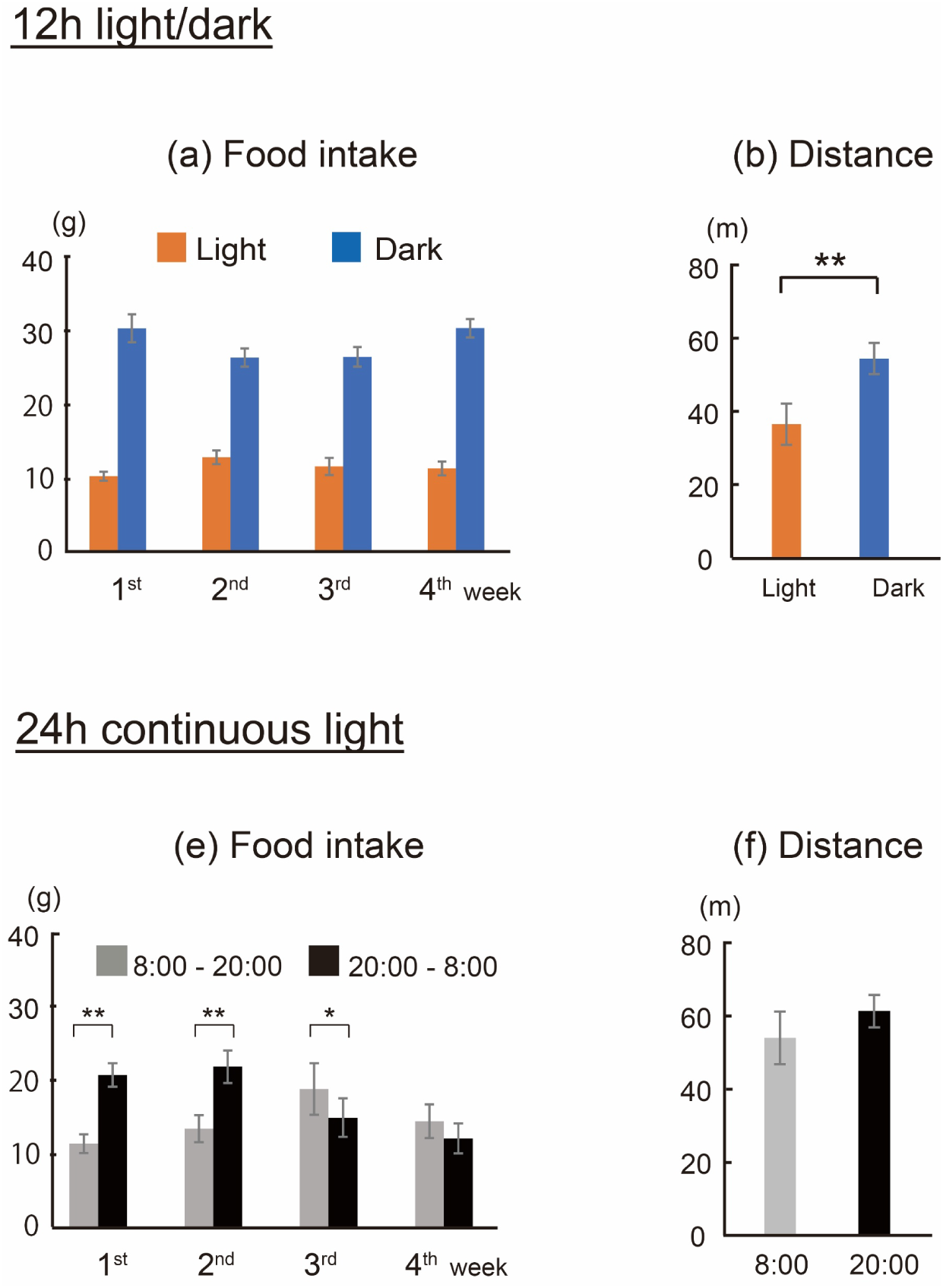
Diurnal changes in food intake and total distance under different light conditions. **(a)** Food intake over 12 h per cage (housing two rats) under 12/12-h light/dark conditions. **(c)** Food intake over 12 h per cage after starting continuous light exposure. **(b, d)** Total locomotor distance per rat over 12 h in the fourth week of continuous light exposure. Paired t-tests were conducted. Under 12/12-h light/dark conditions, food intake and distance traveled during the dark phase were significantly higher than during the light phase (n = 16, p < .01). Under 24-h light exposure, dark-phase food intake was significantly higher than light-phase intake until the third week (p < .05) but no significant difference was observed in the fourth week (n = 16). Total locomotor distance did not differ significantly between the light and dark phases. Error bars represent mean ± SEM. **P < .01.

**Extended Data Fig. 3.**
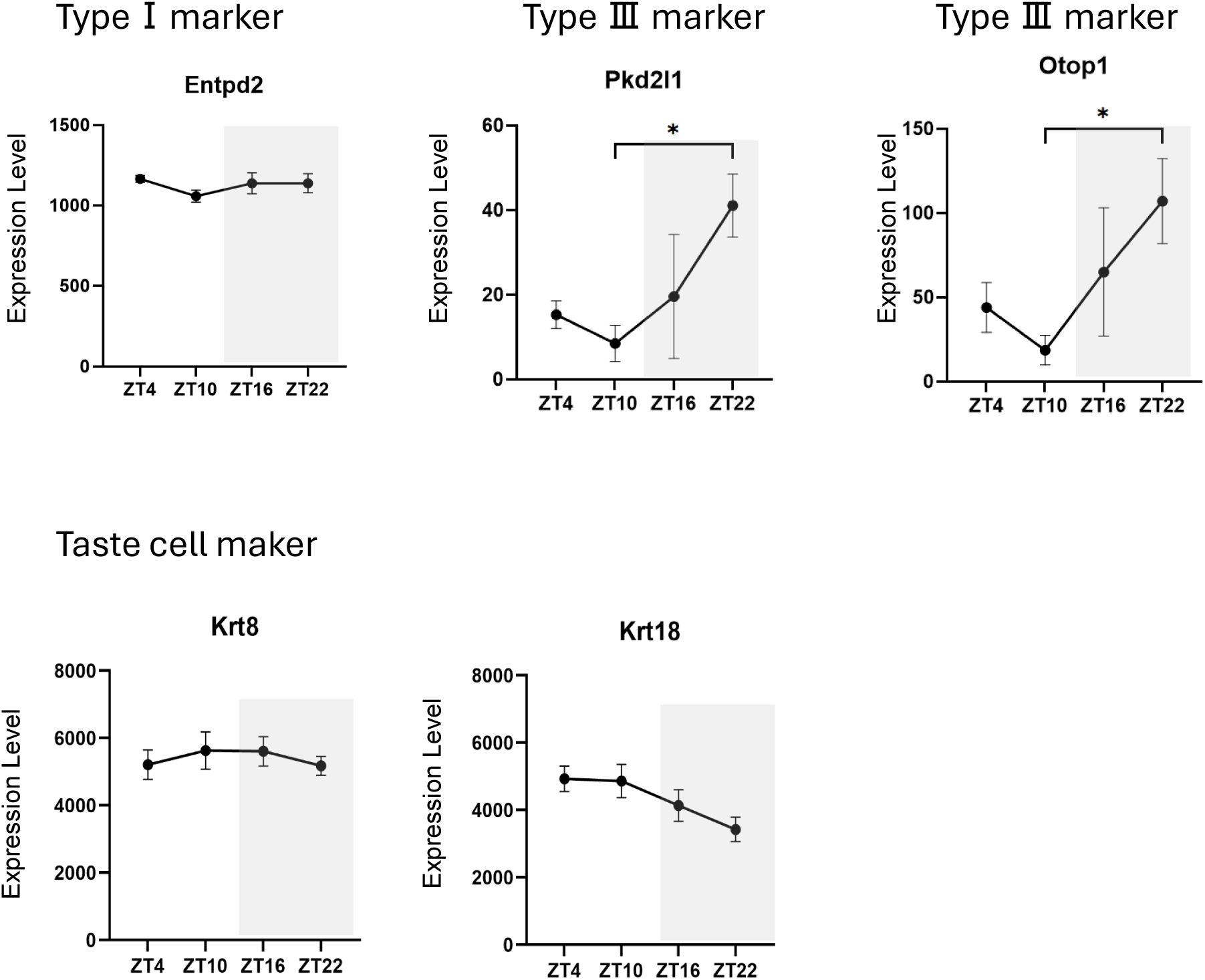
Daily fluctuations in the expression of maker genes for type Ⅰ and type Ⅲ taste cells, along with pan taste cell makers. Kruskal–Wallis test with Dunn’s multiple comparison was applied (n = 6 per condition). Error bars represent mean ± SEM. *P < .05.

**Extended Data Table 1.** Daytime Fluctuations in Physiological and Subjective Indicators in humans.

|  |  | Morning | Noon | Evening |
| --- | --- | --- | --- | --- |
| Body temperature (°C) | Mean | 36.2 | 36.4<sup>a*</sup> | 36.4<sup>b*</sup> |
|  | ±SD | 0.32 | 0.30 | 0.28 |
| Saliva volume (g) | Mean | 0.71 | 0.69 | 0.84 |
|  | ±SD | 0.41 | 0.46 | 0.47 |
| Hunger Score | Mean | 21.8 | 58.8<sup>a**</sup> | 50.7<sup>b**</sup> |
|  | ±SD | 21.3 | 21.8 | 25.0 |
| Fatigue Score | Mean | 2.8 | 2.9 | 4.9<sup>b*,c*</sup> |
|  | ±SD | 3.1 | 3.5 | 4.0 |

One-way ANOVAs with Shaffer post hoc test were conducted, n = 19, respectively.
a; Morning < Noon, b; Morning < Evening, c; Noon < Evening
\*\*p < .01, \* p < .05

**Extended Data Table 2.** Fluctuations in blood glucose and appetite-related hormone concentrations in rats during a day.

|  |  |  |  |
| --- | --- | --- | --- |
|  |  | Light | Dark |
| Glucose (mg/dL) | Mean | 159.2 | 153.5 |
|  | ±SD | 42.0 | 6.6 |
| Leptin (ng/mL) | Mean | 159.0 | 130.9 |
|  | ±SD | 40.4 | 31.8 |
| Ghrelin (pg/mL) | Mean | 719.4 | 717.4 |
|  | ±SD | 77.9 | 258.6 |

T-tests were performed, but no significant differences were detected between light and dark periods in any of the indexes. n = 7, respectively. The Ghrelin values are the sum of acylated and non-acylated ghrelin.

